# Beyond the SNP threshold: identifying outbreak clusters using inferred transmissions

**DOI:** 10.1101/319707

**Authors:** James Stimson, Jennifer Gardy, Barun Mathema, Valeriu Crudu, Ted Cohen, Caroline Colijn

## Abstract

Whole genome sequencing (WGS) is increasingly used to aid the understanding of pathogen transmission. A first step in analysing WGS data is usually to define “transmission clusters”, sets of cases that are potentially linked by direct transmission. This is often done by including two cases in the same cluster if they are separated by fewer SNPs than a specified threshold. However, there is little agreement as to what an appropriate threshold should be. We propose a probabilistic alternative, suggesting that the key inferential target for transmission clusters is the number of transmissions separating cases. We characterise this by combining the number of SNP differences and the length of time over which those differences have accumulated, using information about case timing, molecular clock and transmission processes. Our framework has the advantage of allowing for variable mutation rates across the genome and can incorporate other epidemiological data. We use two tuberculosis studies to illustrate the impact our approach: with British Columbia data by using spatial divisions; with Republic of Moldova data by incorporating antibiotic resistance. Simulation results indicate that our transmission-based method is better at identifying direct transmissions than a SNP threshold, with dissimilarity between clusterings of on average 0.27 bits compared to 0.37 bits for the SNP threshold method and 0.84 bits for randomly permuted data. These results show that it is likely to outperform the SNP threshold where clock rates are variable and sample collection times are spread out. We implement the method in the R package `transcluster`.

## Introduction

Whole genome sequencing (WGS) of pathogens has become an essential tool for improving understanding of how infectious diseases spread between hosts, particularly in the case of tuberculosis (TB) (Hatherell *et al.* 2016). The phylogeny derived from pathogen genomic data helps us to infer likely transmission events. Typically, samples are taken from patients in the field, the date and other epidemiological data are recorded, and the pathogen’s genome is sequenced. A first step is typically to assign cases to clusters; for infectious diseases, a cluster is a group of closely related infections that is usually interpreted as resulting from recent transmission (Poon 2016). These clusters are chosen primarily with the aim of making meaningful subdivisions of the data, with the added benefit of making the amount of data fed into attempts to reconstruct outbreaks and to transmission inference models more tractable. However the assignation method is often somewhat *ad hoc*.

The simplest way to determine sequence relatedness is to count the number of single nucleotide polymorphisms (SNPs) that differ between two sequences. The SNP threshold approach places two individuals in the same putative transmission cluster if there are fewer than a threshold number of SNPs between their sequenced pathogen genomes. Many existing methods to identify outbreak clusters rely on SNP thresholds, as surveyed recently (Hatherell *et al.* 2016) in the case of TB. Similar methods are also used for other pathogens (Dallman *et al.* 2015; Octavia *et al.* 2015). However, there is little agreement in the literature as to what such a threshold should be – see Table 1 for TB SNP thresholds used in some recent studies. The contexts in which these thresholds are applied differ from study to study, so these numbers are not always strictly comparable, but they do indicate the wide range of values that can reasonably be adopted when determining whether or not cases are closely related. By itself, the number of SNP differences between genomes does not directly imply a probability of recent transmission. This is implicitly recognised in some sources. For example, we have from Walker *et al.* (2013): "We predicted that the maximum number of genetic changes at 3 years would be five SNPs and at 10 years would be ten SNPs". Indeed, other studies directly question the use of SNP thresholds, such as Guerra-Assunção *et al.* (2015), Bergholz *et al.* (2014) in the context of food-borne pathogens, and Azarian *et al.* (2016) in an analysis of the spread of methicillin-resistant Staphylococcus aureus (MRSA). Nevertheless, the use of a single SNP threshold is often employed in practice; for example the 12 SNP threshold, used for inferring likely transmission between a pair of TB cases by Public Health England (Walker *et al.* 2014) amongst others, is perhaps the most common in TB.

**Table 1:** SNP thresholds used in recent TB studies. The lower threshold indicates the number of SNPs below which cases are positively identified as belonging to the same cluster. Where different, the upper threshold indicates the the number of SNPs above which cases are identified as clearly not belonging together. Unless otherwise stated, intermediate values are indeterminate.

| Authors | Lower SNP Threshold | Upper SNP Threshold |
| --- | --- | --- |
| Bryant *et al.* (2013b) | $\leq 6$ (relapse) | $> 1000$ (re-infection) |
| Clark *et al.* (2013) | $< 50$ | $> 50$ |
| Guerra-Assunção *et al.* (2015) | $\leq 10$ (relapse) | $> 100$ (re-infection) |
| Lee *et al.* (2015) | $< 2$ | (not specified) |
| Roetzer *et al.* (2013) | $\leq 3$ | (not specified) |
| Walker *et al.* (2013) | $\leq 5$ | $> 12$ |
| Yang *et al.* (2017) | $\leq 12$ | (not specified) |

The appropriate SNP cut-off for inferring transmission is likely to depend critically on the context. There are many sources of uncertainty. Nucleotide mutation rates vary between pathogens, can vary at different stages of infection, and are subject to the effects of selection pressure. Culture processes (e.g. liquid vs solid culture, single colony picks vs sweeps) may affect the diversity in samples that are sent for sequencing. Furthermore, the process of producing finalised SNP data from patient-derived biological samples is a multi-stage procedure where there are choices to be made - including how stringently quality filtering is applied to raw genomic data - which will in general result in different SNP differences being reported. As such, it is important that during every step of the pipeline from sampling from patients and processing the data, to building the models and drawing conclusions from them, that we are aware of sources of uncertainty and attempt to propagate this uncertainty to any conclusions. It is also important that as WGS is rolled out widely as a tool in infectious disease, we re-calibrate SNP-based methods to accommodate changes in both sequencing technologies and in the bioinformatics pipelines used to call variant SNPs. Clustering methods that use variant SNP calls exclusively will be most sensitive to such changes.

The fundamental logic behind SNP cut-offs is that it takes time to accrue genetic variation; even in organisms where the molecular clock is variable, it seems uncontroversial to assume that two isolates that differ by only a few SNPs are more likely to be a result of recent transmission than isolates that are 50 SNPs apart. However, the rate at which polymorphisms occur varies not only between organisms (Kuo and Ochman 2009), but also across a genome; it is affected by selection pressure and by horizontal gene transfer (HGT) (Novichkov *et al.* 2004), though this is not an issue for TB and there are methods to remove recombination and HGT prior to using SNP cut-offs. As per Barrick and Lenski (2013), it is also important to distinguish between the *mutation rate*, the rate at which spontaneous mutations occur, and the *substitution rate*, the rate of accumulation of changes in a lineage; this depends on both the mutation rate and the effects of selection and drift. Here, when we refer to the clock rate, we mean the substitution rate, as we use the rate to interpret variants measured with sequencing technologies.

This distinction is particularly important for diseases like TB, where selection pressure due to antibiotics can be substantial. Whilst the background SNP accumulation rate for *Mycobacterium tuberculosis* has been estimated at 0.5 SNPs/genome/year (Walker *et al.* 2013), selection pressure and antibiotic resistance can influence this rate considerably. For example, in Eldholm *et al.* (2014) we see the observation that “After exclusion of transient mutations in the patient isolates, 4.3 mutations were acquired per year … or 2.3 mutations per year when excluding resistance mutations.” The size of the population of bacteria within a host could also affect the number of SNPs observed between that host and those they infect. Unexplained larger variation is also encountered as documented in Korhonen *et al.* (2016), though high SNP numbers could be a result of re-infection or mixed infection rather than in-host evolution. Where we know that selection or high substitution rates are likely to be present and detected, a higher rate is therefore likely to be appropriate for clustering, and this will affect the relationship between SNPs and transmission events.

It should be noted that there are other approaches to clustering, based on molecular (but not WGS) data and including time and geographical data. For example, Kammerer *et al.* (2013) apply three different statistical tools to spoligo-type data and mycobacterial interspersed repetitive units (MIRU) data, together with date and location of cases, to show that these tools can successfully identify TB outbreaks. Donker *et al.* (2016) use variable number tandem repeat (VNTR) data for MRSA cases, with time and location data, to identify clusters based on a hierarchical clustering method (Ypma *et al.* 2013). There are also software packages such as *vimes* (Jombart and Cori 2017), which provides tools allowing users to integrate different types of data and detect outbreaks. These approaches treat each of the underlying variables as independent inputs, without explicitly modelling the connection between time and the accumulation of genetic differences.

We take a slightly different tack here, and jointly use the sample time and genetic distance, together with a model of SNP acquisition over time and transmission events over time, to base putative transmission clusters on the probability that cases are separated by a threshold number of transmission events. This is motivated by a belief that the number of transmission events between two cases is a natural and intuitive measure of how "clustered" they are in the sense of transmission (and how likely they are to be part of the same outbreak). This cannot usually be measured directly and must be inferred from other data. However, it is reasonable to assume that appropriate incorporation of the time over which the accumulation of SNPs occurs, as well as the likely time between transmission events, give a more accurate and nuanced measure of the likelihood that cases are linked by a small number of transmission events. We develop a probabilistic approach which permits variation in the SNP accumulation process, allows for faster SNP accumulation for sites under selection and allows for variation in the speed with which individuals infect their contacts. We aim to provide a principled alternative to SNP cut-offs for clustering pathogen genomes into putative transmission clusters.

## New Approaches

Two samples are usually considered to be in the same transmission cluster if the number of SNPs between them is less than or equal to a fixed cut-off, or threshold. This is a quick way to explore relatedness among a group of isolates and gain an approximate understanding of the extent of recent (low-distance) transmission, but it is coarse and embeds a number of strong assumptions.

Our proposed probabilistic transmission approach, in contrast, is based on sample pairs being clustered together if we estimate that there were fewer than a threshold number of transmission events between them, with a given probability. It uses the same genetic (SNP) distance information as the SNP threshold method, but in addition makes use of the sample times, knowledge of the SNP accumulation and transmission processes. The essential inputs to our method are: the number of SNP differences between sample pairs, the sample dates, the assumed clock rate and the assumed transmission rate.

In addition, our method can readily be extended to incorporate other factors: we show in Materials and Methods how this can be done for spatial data, and for antibiotic resistance. Building these inputs into our model allows us to create a more nuanced and principled way of identifying transmission clusters, and allows us to apply the method consistently in varied settings - for example, in those where drug resistance is suspected to be a factor.

We start by establishing probability distributions for the total length of time (*h* years) along both lines of descent from the most recent common ancestor (MRCA) of a pair of samples; this depends on the clock process, and helps define the distance between the two samples (there is *h*/2 years of elapsed time from the MRCA to the earlier sampling date). We then compute the probability that at least a threshold number of transmissions took place between the two sampled cases over this time, where the probability distribution for the number of transmissions *k* is *P*(*k*|*h*) (see Table 6 for a summary of the symbols used and their units). This approach gives the flexibility to incorporate sample time information and other data. The method uses sample dates and aligned sequence data (variant calls) together with models of the clock and transmission processes. For a pair of samples, we use the SNP distance *N*, the time difference between their sampling dates (*δ* years) and the clock process to write down the probability distribution 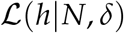 for when the most recent common ancestor of the two sequences existed; this must be before the first sampled case. Integrating over this unknown time, we can find the probability that a certain number of transmissions separate the two cases:

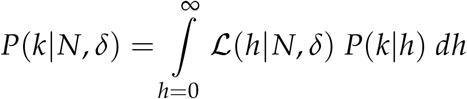

This is Equation (9), developed in more detail in Materials and Methods. To incorporate spatial data, a weighting *w* is applied to the probabilities to reflect that spatial distance can affect estimates of the number of intermediate transmissions between two sampled individuals; we express this in Equation (19):

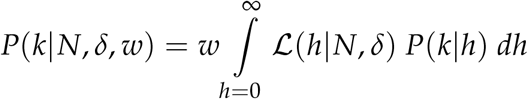

## Results

We illustrate how the transmission method compares to the SNP threshold method for a simple toy example. We define the “T cut-off” as the cut-off level for the transmission method, using Equation (10); the samples are clustered together where the implied number of transmissions *k* is less than or equal to T with a probability of 80%, given some clock rate *λ* and transmission rate *β*. We see that the transmission method clusters the cases together in a different order to the SNP threshold method as the cut-off level is incremented. Cases A and B are the closest in SNP distance, but the time elapsed between their sampling dates increases their distance by the transmission distance function relative to cases C and D, which are sampled at the same time as each other. So when we take timing into account, the clustering is altered (also illustrated in Figure 1).

**Figure 1:**
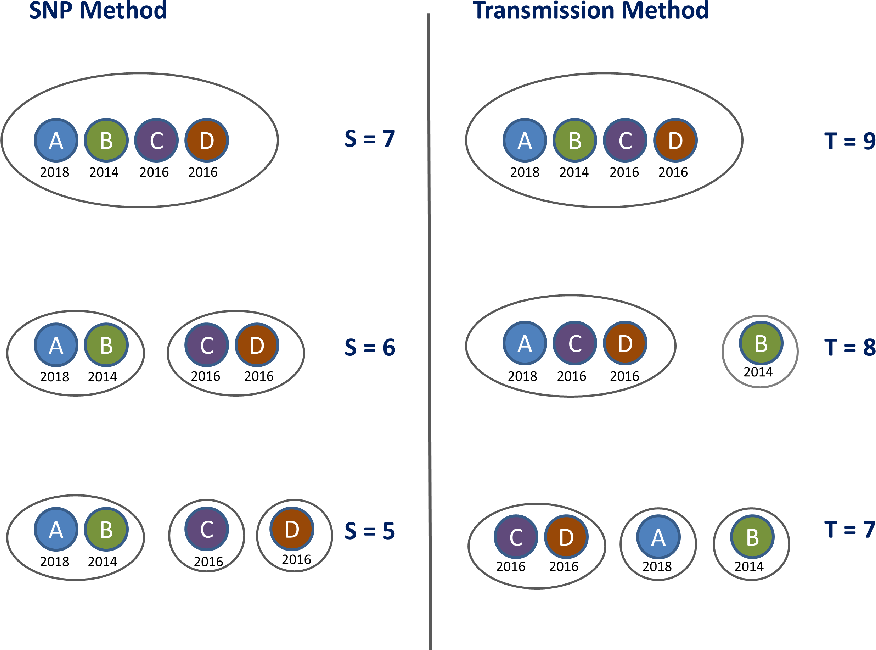
Clustering on the toy example data set provided in Table 2. The left hand panel shows the clusters obtained by applying the SNP threshold method with three different thresholds, with the cut-off level denoted by S; samples are clustered together where the SNP distance is less than or equal to S. The right hand panel shows the clustering obtained by applying the transmission method, using Equation (10), with the cut-off level denoted by T; samples are clustered together where the implied number of transmissions *k* is less than or equal to T with a probability of 80%, with clock rate *λ* = 1.5 SNPs/genome/year and *β* = 2.3 transmissions/year.

We model the number of intermediate transmissions between two sampled hosts given the total time over which SNPs have likely accumulated. Altering the transmission rate *β* (by which we mean the rate at which intermediate cases occur in the total time elapsed *between the MRCA of two sampled hosts and the sampling events*; see Materials and Methods) alters the absolute transmission cut-off level at which the clusters change - in this example, increasing *β* to 3.0 transmissions/year gives the same clusters as in Figure 1 but at levels 9, 10 and 12 transmissions rather than 7, 8 and 9 transmissions respectively. This has no impact on the order of the clustering as the level of the cut-off changes.

**Table 2:**
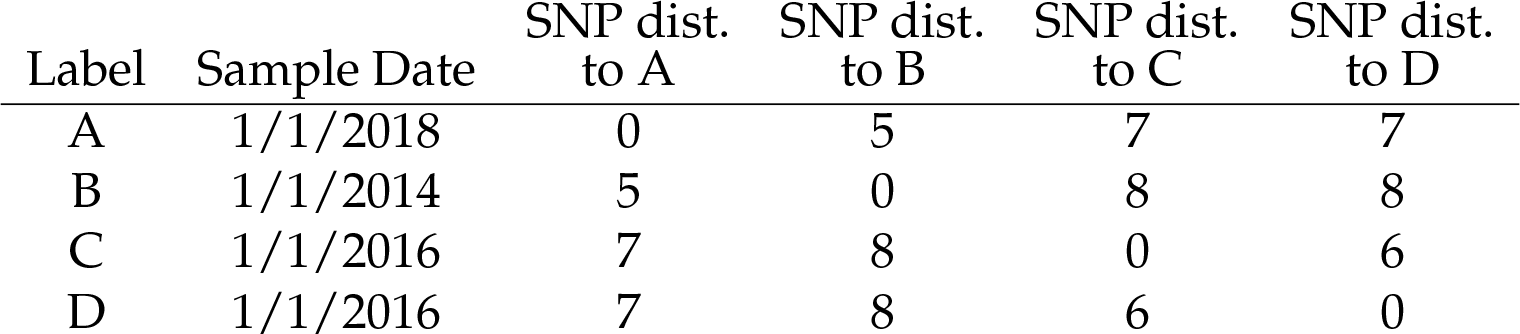
Model inputs for toy example data set.

| Label | Sample Date | SNP dist.<br>to A | SNP dist.<br>to B | SNP dist.<br>to C | SNP dist.<br>to D |
| --- | --- | --- | --- | --- | --- |
| A | 1/1/2018 | 0 | 5 | 7 | 7 |
| B | 1/1/2014 | 5 | 0 | 8 | 8 |
| C | 1/1/2016 | 7 | 8 | 0 | 6 |
| D | 1/1/2016 | 7 | 8 | 6 | 0 |

By contrast, altering the clock rate does have a material impact on the way clustering occurs as we increase the transmission threshold. For *λ* = 0.5 SNPs/genome/year, cases A and B are closest under the transmission method, just as they are with the SNP threshold method, and so the clustering is the same for both methods. At *λ* = 1.5 SNPs/genome/year, cases C and D are closest under the transmission method, and the clustering evolves as shown in Figure 1.

### British Columbia data

We analyse a data set from British Columbia, comparing the SNP threshold method to the transmission method in Equation (10). The data set comprises 52 samples collected from 51 patients over a 14 year period, and has been pre-filtered with the result that all samples are relatively close - within 25 SNPs. Consequently, using the SNP threshold method with the threshold set to 13 SNPs or higher, all samples are placed in one cluster. When the threshold is 9 SNPs, we obtain a large 42-case cluster, a secondary 8-case cluster and some outliers. As we reduce the threshold further down to 3, the large cluster breaks up but the 8-case cluster persists. We illustrate this in Figure 2.

**Figure 2:**
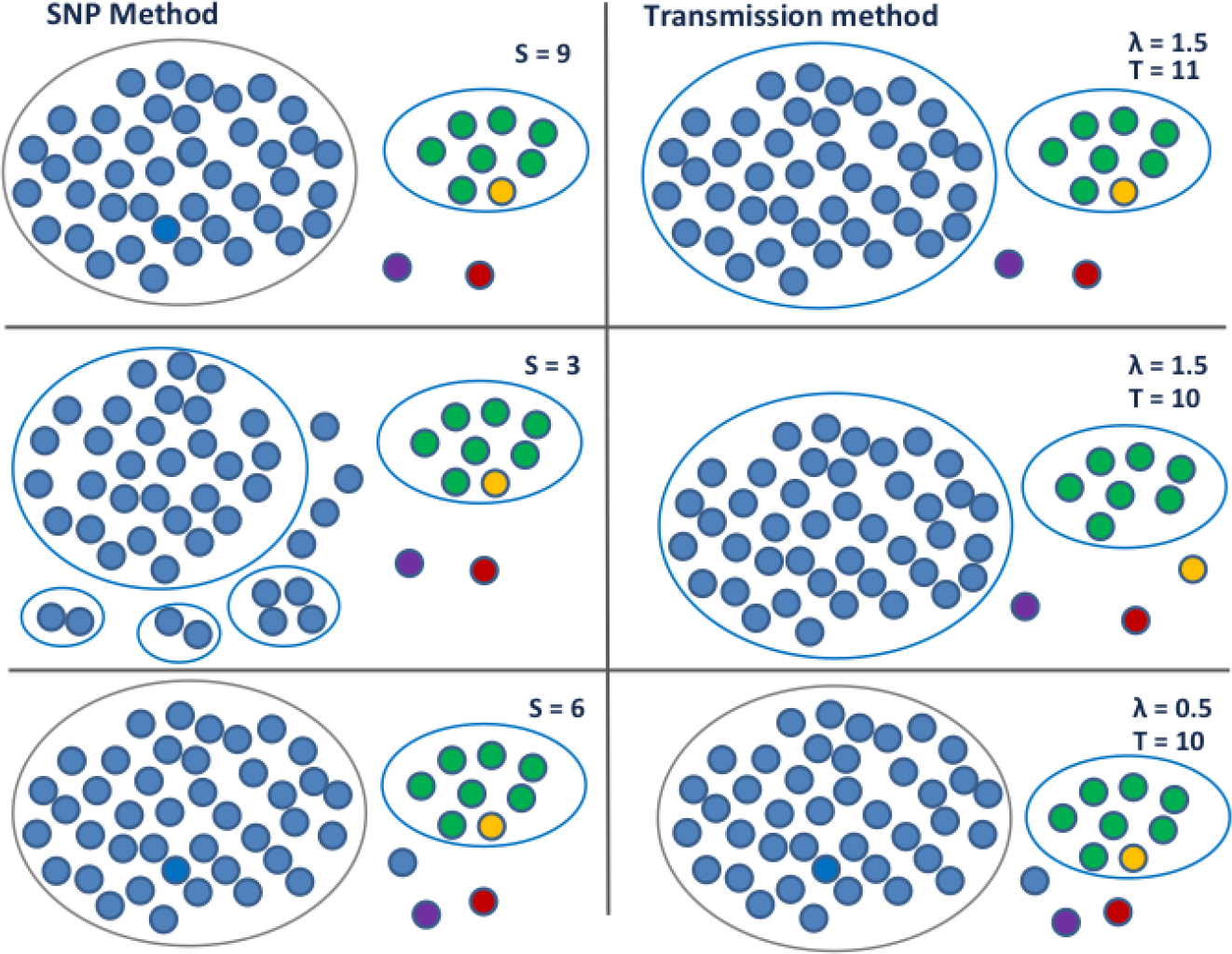
Clustering on the British Columbia data set. The left hand side shows the clusters obtained by applying the SNP threshold method with three different thresholds, with the cut-off level denoted by S. The largest cluster breaks up as the level is lowered whilst the size 8 cluster remains intact. The right hand side shows the clustering obtained by applying the transmission method, using Equation (10); samples are clustered together where the implied number of transmissions *k* is less than or equal to T with a probability of 80%. As shown in the top two thirds, with clock rate *λ* = 1.5 SNPs/genome/year and *β* = 2.0 transmissions/year, the size 8 cluster loses a member whilst the largest cluster stays the same as the level is lowered. When *λ* is low, *h* is larger, so the MRCA of a cluster gets pushed back further in time. In this case, the value of *δ* between two cases has a limited impact on the estimated number of transmissions; the SNP difference is dominant, and we recover the same clustering that is obtained with the SNP cut-off. This is shown in the lower third, where the clock rate *λ* = 0.5 SNPs/genome/year and *β* = 1.2 transmissions/year.

We used the transmission method, using Equation (10) with *β* = 2.0 transmissions/year and two different average clock rates: *λ* = 0.5 and 1.5 SNPs/genome/year (1.5 is larger than the typical rate for TB but within other outbreak estimates (Bryant *et al.* 2013a)). When *λ* is low we can obtain the same clustering as with the SNP cut-off. When *λ* is higher, we have one cluster which contains all the samples for *T* > 11. As with the SNP threshold method, at *T* = 11 we have a large 42 case cluster, a secondary 8-case cluster and some outliers. But as we move to *T* = 10, the secondary cluster loses a member, whilst the main cluster stays at size 42. This is because one of the members of the secondary group is very close by the SNP distance to another member of that group, but was sampled more than 10 years before. As with our simple toy example, timing alters the effective distance between samples because the distance into account the clock rate and the transmission rate, and so the timing information can affect the clustering.

Furthermore, the probabilistic nature of the approach means that we can see how strongly we predict cases to be linked; in Figure 3 we use thicker edges to denote a higher probability of being linked by relatively few transmissions. In addition, we show the effect of incorporating spatial proximity, using Equation (19); we assign each of the cases into one of six numbered regions. Including a spatial weighting, reflecting the barrier to the infection moving between different regions, and leaving all other parameters unaltered, changes the clustering that is obtained.

**Figure 3:**
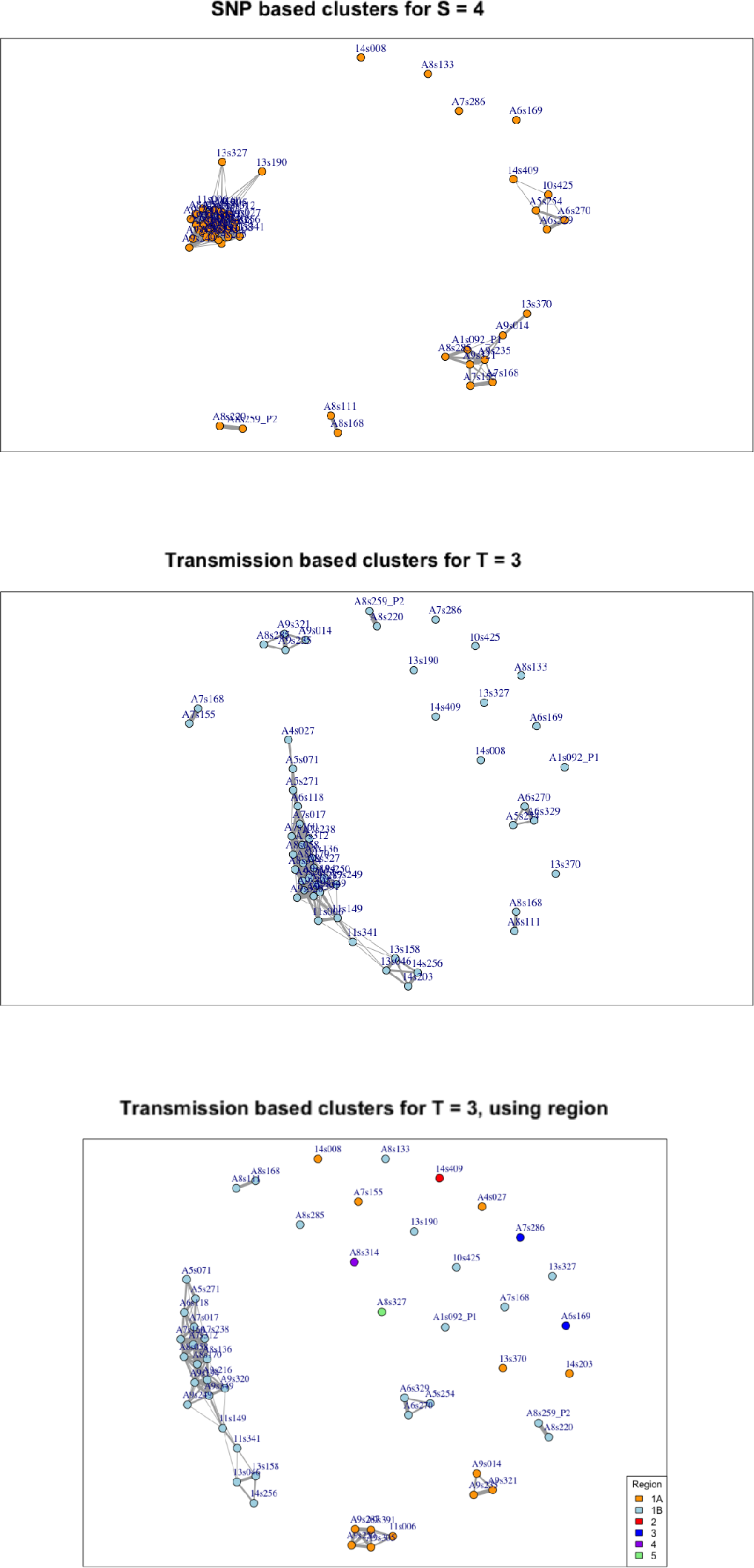
Three views of the same British Columbia TB data illustrating the contrasting effect of implementing the SNP and transmission methods and showing estimates of how close individual cases are to each other. In the top figure, edges between nodes indicate that cases are within 4 SNPs of each other. In the lower figures, edges indicate that cases are 80% likely to be within 3 transmission events of each other, given a clock rate *λ* = 1.5 SNPs/genome/year and *β* = 2.0 transmissions/year. The middle figure is based on Equation (10), and the bottom figure uses Equation (19), with weighting *w* = 20% where two cases are assigned to differing regions. The thicker the edges, the closer the cases are: for the SNP based clusters the thickest edges correspond to no SNP difference, the thinnest to a distance 4 SNPs; for the transmission based clusters the thickest edges correspond to one likely transmission event, the thinnest to 3.

#### Sensitivity to clock rate

An implicit assumption of the SNP threshold method is that each SNP contributes equally towards the SNP distance. This implies that the clock rate or substitution process is constant across the set of isolates and across the genome. When the same threshold is used in different settings and across different pathogen sub-types, the implicit assumption is that the same substitution process holds in these settings. In our new transmission method, the effective distance between any two samples is inversely proportional to the assumed mean clock rate. A lower clock rate means that more time is needed in order for a fixed number of SNPs to be generated; this gives room for more potential intermediate transmission events. A higher clock rate means that the fixed time between samples has a greater effect on the clustering, as the time between samples places a greater constraint on the range of possible heights *h*; the fixed time "uses up" more of the time available than it would under a low clock rate (because there is less total estimated time available, a higher portion of it is in the time period *δ*). We show this in Table 3: the transmission clustering method approaches the same results as the SNP clustering method as the assumed clock rate is reduced. We use the variation of information dissimilarity measure given by *clue* (Meilă 2007) to compare the clusters produced by the two methods.

**Table 3:** Effect of varying the clock rate using the British Columbia data. This table shows how the clock rate affects the transmission method, using Equation (10), keeping the transmission rate *β* constant. For the SNP threshold method, samples are clustered together where the SNP distance is less than or equal to S. For the transmission method, samples are clustered together where the implied number of transmissions *k* is less than or equal to T with a probability of 80%. For a clock rate of 0.5 SNPs/genome/year, the transmission method matches all SNP-based clusters for thresholds of 5 SNPs and above. As the clock rate increases, the transmission clustering diverges further from the SNP clustering. As we vary the other parameters, the choice of *β* is effectively a scale factor and does not affect the pattern of clustering. We use the variation of information dissimilarity measure given by *clue* (Meilă 2007) to compare the results of the two methods.

| SNP Threshold | Largest cluster | $\lambda$ | $\beta$ | Trans. Threshold | Largest cluster | Dissimilarity between SNP and Trans. methods |
| --- | --- | --- | --- | --- | --- | --- |
| S=12 | 50 | 0.5 | 1.2 | T=22 | 50 | 0.000 |
| S=9,10,11 | 42 | 0.5 | 1.2 | T=16 | 42 | 0.000 |
| S=6,7,8 | 41 | 0.5 | 1.2 | T=10 | 41 | 0.000 |
| S=5 | 37 | 0.5 | 1.2 | T=8 | 37 | 0.000 |
| S=4 | 31 | 0.5 | 1.2 | T=7 | 31 | 0.139 |
| S=3 | 29 | 0.5 | 1.2 | T=6 | 29 | 0.052 |
| S=2 | 28 | 0.5 | 1.2 | T=3 | 28 | 0.113 |
| S=12 | 50 | 1.0 | 1.2 | T=11 | 50 | 0.000 |
| S=9,10,11 | 42 | 1.0 | 1.2 | T=8 | 42 | 0.000 |
| S=6,7,8 | 41 | 1.0 | 1.2 | T=5 | 41 | 0.058 |
| S=5 | 37 | 1.0 | 1.2 | T=4 | 37 | 0.113 |
| S=4 | 31 | 1.0 | 1.2 | T=3 | 29 | 0.374 |
| S=3 | 29 | 1.0 | 1.2 | T=3 | 29 | 0.174 |
| S=2 | 28 | 1.0 | 1.2 | T=1 | 28 | 0.113 |
| S=12 | 50 | 1.5 | 1.2 | T=7 | 52 | 0.163 |
| S=9,10,11 | 42 | 1.5 | 1.2 | T=6 | 42 | 0.000 |
| S=6,7,8 | 41 | 1.5 | 1.2 | T=4 | 41 | 0.058 |
| S=5 | 37 | 1.5 | 1.2 | T=2 | 35 | 0.289 |
| S=4 | 31 | 1.5 | 1.2 | T=2 | 35 | 0.400 |
| S=3 | 29 | 1.5 | 1.2 | T=1 | 29 | 0.240 |
| S=2 | 28 | 1.5 | 1.2 | T=1 | 29 | 0.290 |
| S=12 | 50 | 2.0 | 1.2 | T=6 | 52 | 0.163 |
| S=9,10,11 | 42 | 2.0 | 1.2 | T=4 | 42 | 0.058 |
| S=6,7,8 | 41 | 2.0 | 1.2 | T=4 | 42 | 0.149 |
| S=5 | 37 | 2.0 | 1.2 | T=2 | 37 | 0.412 |
| S=4 | 31 | 2.0 | 1.2 | T=1 | 29 | 0.447 |
| S=3 | 29 | 2.0 | 1.2 | T=1 | 29 | 0.230 |
| S=2 | 28 | 2.0 | 1.2 | T=1 | 29 | 0.240 |

### Moldova data

This data set comprises 422 samples collected over a period of less than 2 years. For this data - with any reasonable choices of parameters and a fixed substitution rate for all sites - as shown in Table 4, our new transmission method does not differ from the SNP threshold method. This can be explained by two factors that work together: the small distance in time between any two samples and the large SNP differences between cases. There isn’t enough variation in the timing information relative to the SNP distances for an appreciable difference to emerge between the two clustering methods.

**Table 4:**
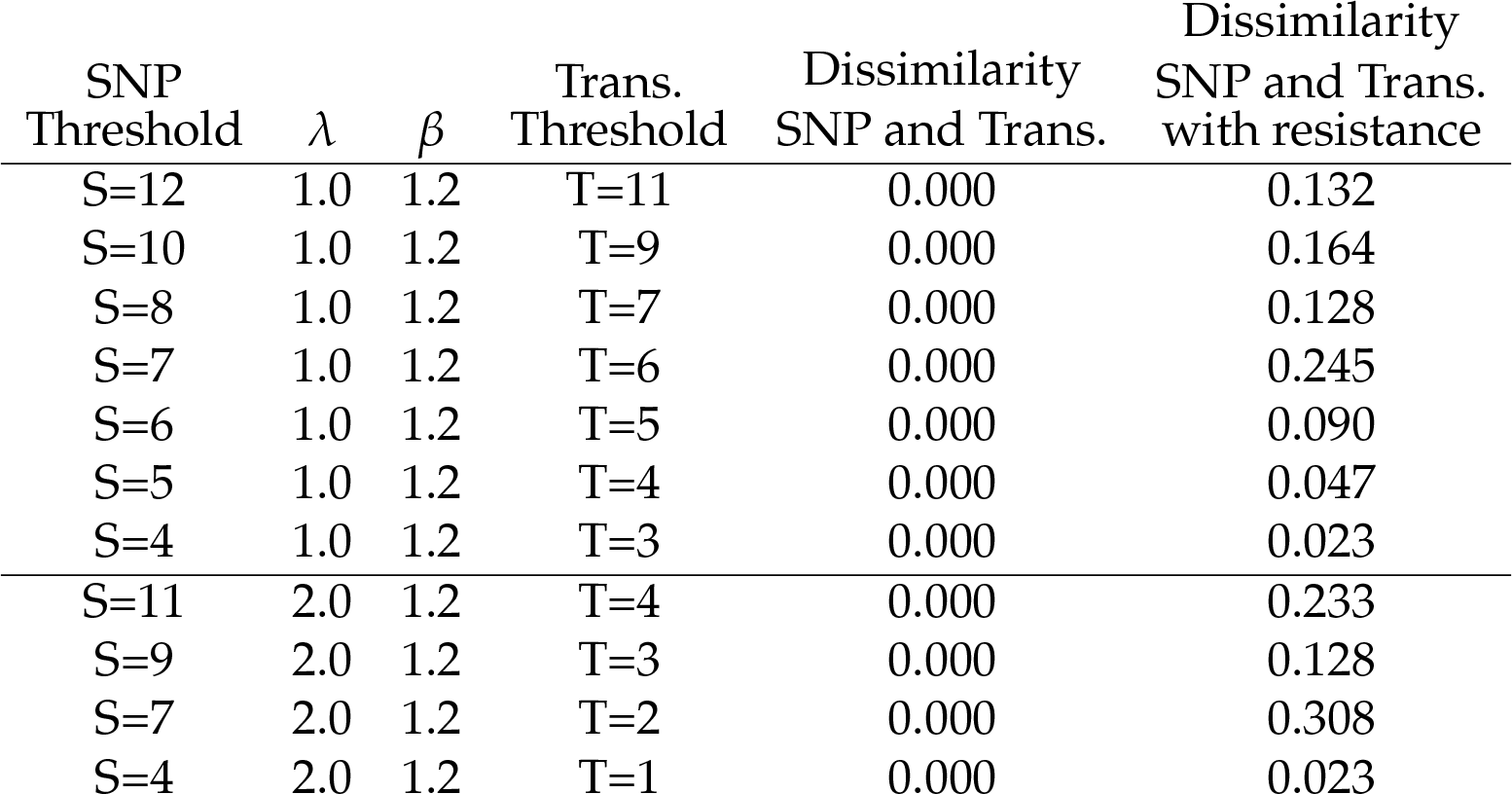
Comparison of methods with the Moldova data, taking drug resistance into account. This table shows how the clock rate affects the transmission method, using Equation (10), keeping the transmission rate *β* constant, and the effect of including resistance using Equation (18). For *λ* = 1.0 and 2.0 SNPs/genome/year, the pattern of clusters is identical using the SNP and transmission methods across a range of thresholds, but differs when resistance-conferring SNPs are taken into account. For the SNP threshold method, samples are clustered together where the SNP distance is less than or equal to S. For the transmission method, samples are clustered together where the implied number of transmissions *k* is less than or equal to T with a probability of 80%. We use the variation of information dissimilarity measure given by *clue* (Meilă 2007) to compare the results of the methods.

#### Use of drug resistance-conferring SNPs

We can, however, explore the role of drug resistance-conferring SNPs on the clustering. Information on the location of resistance-conferring sites for TB was obtained using PhyResSE (Feuerriegel *et al.* 2015) and a resistance-conferring SNP distance matrix was computed for the Moldova data by filtering against this information. Selection is likely to lead to resistance-conferring SNPs arising more quickly than other SNPs: for example, one TB study (Eldholm *et al.* 2014) gives a mutation rate of 4.3 SNPs per genome per year when they are included, in contrast to the 0.5 SNPs per genome per year that is typically estimated for TB (Walker *et al.* 2013). Resistance acquisition may further increase the rate of acquisition of additional SNPs through multiple resistance, compensatory mutations or other mechanisms. For this analysis we used a clock rate for the drug-resistant sites, as in Equation (18), which is five times higher than for the sites which are not resistance-conferring.

Overall, resistance-conferring SNPs in the Moldova data set form only 0.6% of the total number of SNPs. However, restricting to those sample pairs where the SNP distance is less than or equal to 20, they form 8% of the total. If a high proportion of the SNPs between two cases are resistance-conferring SNPs, then this effectively shortens the distance between the cases, making them more likely to be joined together in a transmission cluster. For several sample pairs in this data set, the proportion of resistance-conferring SNPs that differ between the two samples is approaching 35%, whilst for some other pairs there are none at all. For this reason we see a difference when we take resistance into consideration, as seen in Table 4 and Figure 4. The largest cluster is not shown in detail in the figure and is more robust with respect to the effect of resistance-conferring SNPs than the smaller clusters.

**Figure 4:**
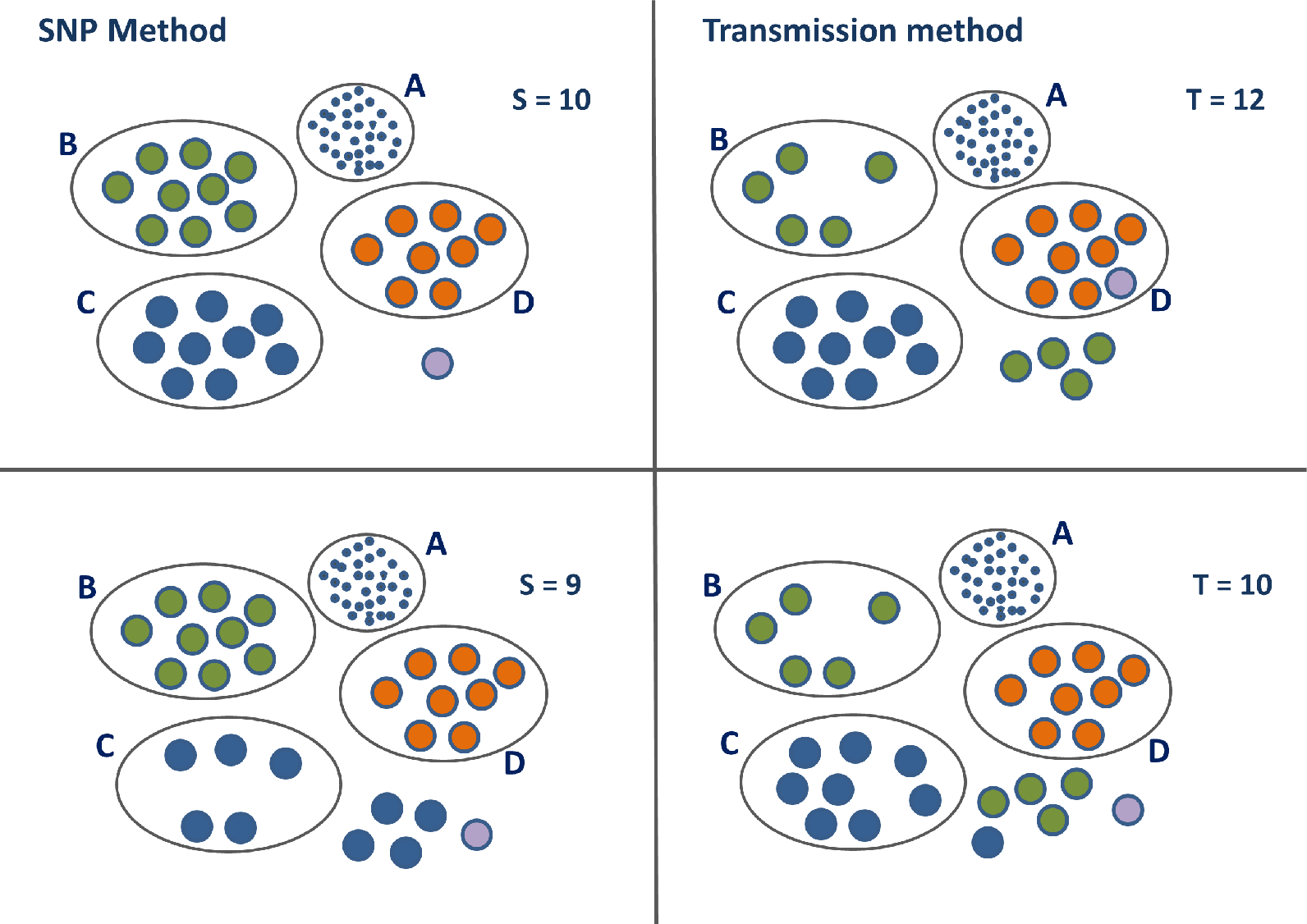
Clusters in the Moldova data set illustrating the effect of accounting for resistance-conferring SNPs in the transmission method, using Equation (18). Clusters B, C and D are the second to fourth largest clusters in the Moldova data using the SNP threshold method. The largest cluster A, with 93 members for *S* = 10 is shown for com-pleteness. Isolated cases are shown with no enclosing oval. Colours are chosen to enable identification of the same cases in the four different scenarios. The left hand panel shows the clusters obtained by applying the SNP threshold method with two different thresh-olds, with the cut-off level denoted by *S*; samples are clustered together where the SNP distance is less than or equal to *S*. The right hand panel shows the clustering obtained by applying the transmission method, using Equation (10), with the cut-off level denoted by T; samples are clustered together where the implied number of transmissions *k* is less than or equal to T with a probability of 80%, with clock rate *λ* = 1.5 SNPs/genome/year and *β* = 2.0 transmissions/year.

### Simulated data

To explore the performance of the clustering methods in a setting where the “ground truth” is known, we simulate data and compare the SNP and transmission (as given by Equation (10)) methods. The "true" clusters are generated from simulated transmission networks produced by *TransPhylo*.

We consider clustering cases based on direct transmission, so that two cases are joined in a cluster if one infected the other, and we compare clusters generated by the SNP threshold method with those generated by the transmission method. In order to compare to the appropriate set of clusters, we find the best match that the method achieves against the true cluster over an appropriately wide range of threshold levels. Then we use the variation of information dissimilarity measure given by *clue* (Meilă 2007) to compare the results of the two methods to the true clusters. We also compare randomly permuted simulated data to the simulated clusters to provide a yardstick of accuracy. This is achieved by fixing the number of clusters to be the number of the true clusters, and then randomly allocating each sample case to one one of those clusters. The results in Table 5 show that the transmission method is consistently better than the SNP threshold method at identifying direct transmissions within an outbreak. Both methods perform significantly better than the randomly generated data.

**Table 5:**
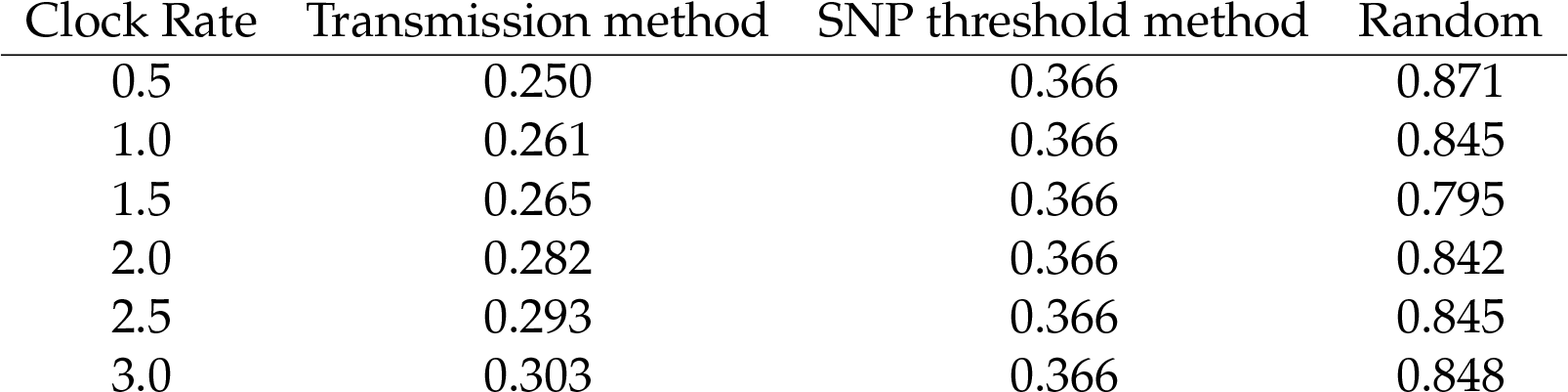
Dissimilarity measure comparing both the SNP and transmission methods against simulated data, averaged over the full set of simulations. We use the variation of information dissimilarity measure given by *clue* (Meilă 2007) to compare the results of the methods. Lower numbers indicate sets of clusters that are more similar to the true clusters. An outbreak was simulated 100 times with 10 sampled cases from a total of between 20 and 30 cases, depending on the simulation. The measure is obtained by comparing clusters from a range of thresholds to the known clusters, and picking the one with the lowest score. The averages are 0.27 bits for the transmission method, 0.366 for the SNP threshold method and 0.84 for the randomly permuted data. The clock rate is the rate used by the transmission method only to relate the number of SNPs to the time distribution, and thus does not affect the SNP threshold method results - this is why the SNP threshold method has the same dissimilarity compared to the simulated data whatever the clock rate. The table shows how the dissimilarity varies as the clock rate varies, for a fixed *β* = 3.0 transmissions/year, as compared to simulated samples connected by direct transmission. The random column shows the dissimilarity obtained for randomly allocated simulated clusters.

| Clock Rate | Transmission method | SNP threshold method | Random |
| --- | --- | --- | --- |
| 0.5 | 0.250 | 0.366 | 0.871 |
| 1.0 | 0.261 | 0.366 | 0.845 |
| 1.5 | 0.265 | 0.366 | 0.795 |
| 2.0 | 0.282 | 0.366 | 0.842 |
| 2.5 | 0.293 | 0.366 | 0.845 |
| 3.0 | 0.303 | 0.366 | 0.848 |

**Table 6:** Symbols used in the model and their meaning.

| Symbol | Meaning | Units |
| --- | --- | --- |
| $\lambda$ | Clock rate | SNPs/genome/year |
| $\beta$ | Transmission rate | Transmissions/year |
| $h$ | Total time over which SNPs occur over both lines of descent before the first sampling date | Years |
| $\delta$ | Time between two sample dates | Years |
| $N$ | No. SNP differences between two cases | |
| $N_\delta$ | No. SNP differences occurring after first sample | |
| $k$ | Number of transmissions | |
| $S$ | Cut-off threshold for SNP threshold method | |
| $T$ | Cut-off threshold for transmission method | |

Identifying direct transmissions is not the aim of either the SNP cut-off or transmission clustering method; rather, both aim to simply group cases into sets of isolates for onward, more intensive (model-specific, Bayesian for example) out-break reconstructions. Testing the ability of SNP vs transmission-based methods to accomplish this using simulated data would require an appropriate simulation set-up, which in turn would have a lot of flexibility (and could no doubt be tweaked to ensure that the transmission method performs well, or that the SNP cut-off does). For example, one approach would to simulate the introduction of new cases whose SNP distance is 25 from existing cases in an existing outbreak. The SNP threshold method with a threshold of 12 SNPs will always correctly place such new introductions in a new cluster, and will group their descending infections correctly until one or more of them is more than 12 SNPs away from other sampled cases in the cluster. Conversely, if new introductions were only 12 SNPs from existing cases the SNP threshold method would mis-classify them as linked to existing clusters. In the transmission method, we can compute the probability that a newly introduced case that is 25 SNPs from existing cases will fall within a certain number of transmission events. This gives us the probability that we would infer an incorrect link to an existing cluster. With *λ* = 1.2 SNPs/genome/year and *β* = 1.5 transmissions/year, the probability that there are more than 10 transmissions for cases 25 SNPs apart is 99.9%. This falls to 98.3% for more than 15 transmissions. Accordingly, the simulation approach for introducing new clusters will greatly affect the performance of both the SNP and transmission-based methods, and so we have not chosen to perform extensive simulations to compare the methods.

## Discussion

We have demonstrated how our approach can be consistently applied in different contexts, with timing information, with spatial data in the case of British Columbia, and with resistance data in the case of Moldova. This is an advance on what is possible with the fixed SNP threshold approach, where there is no general way to adjust thresholds to take this context-specific information into account.

A fixed number of SNPs can arise from different numbers of transmissions depending on other factors, including the timing of transmission, selection for resistance, the substitution process, location and factors we have not explicitly modelled (social contacts, host risk factors, pathogen factors). We have seen that sampled cases which are relatively close in genetic distance can nevertheless be separated by large distances in time. In this scenario, a simple SNP cut-off may place samples too close together for outbreak clustering purposes. In contrast, our new method is robust with respect to outlying cases which have been sampled at very different times compared to the majority of cases. These cases can make inference of timed phylogenetic trees challenging because the low genetic variation is hard to reconcile with the large time distance. Furthermore, true transmission clusters need not be clades in phylogenetic trees, because one cluster could descend from another but be separated by a long time or a large genetic distance (due to sampling effects). Accordingly, the clusters obtained by our method do not necessarily correspond to phylogenetic clades. We briefly discuss the application of our method to timed phylogenetic trees in the supplementary data, with an example cluster which is not a clade.

Our probabilistic transmission method has certain advantages. It is relatively simple, requiring only the implementation of fast-running algorithms to estimate the time distributions; the heavy machinery to run large simulation methodologies (like MCMC) is not required. The amount of information required for the model is limited and consists of as little as the SNP distances, the timing data and a knowledge about the substitution and transmission processes. Nevertheless it has the flexibility to be able to handle SNPs under selection, SNPs with a different substitution process and variability in the substitution and transmission processes, and it has the scope for extensions to include more epidemiological data. Even in data sets where there is not much timing information to work with, we have seen that the integration of information on resistance-conferring sites can be used within our framework to fine tune the clustering. Using two distinct processes - transmission, and the accumulation of measurable genetic variation – to define clusters carries the advantage that these processes may be estimable from data. This enables transmission clusters to be formed based on focused discussion and estimation of measurable processes rather than based on fixed cutoffs, and it allows ready adaptation for new pipelines that detect variation.

There are some limitations. Prior knowledge of the substitution and transmission processes is required, and there is some uncertainty in choosing appropriate values. However, the model is typically robust with respect to changes in these variables; in particular, varying the transmission rate does not have a material impact on the clustering because a re-scaling of the cut-off will compensate. The choice of a time-varying transmission function *β*(*t*) is, however, likely to have an impact on results. In particular we would expect a low probability of very quick transmission – as the pathogen numbers are building up in a new host – to have a significant impact, compared to the use of a constant transmission rate, as would a fast rate early diminishing to a much lower rate later. Note also that the parameter *t* in our model represents the total time since infection to both the sample dates: so we are not modelling the variation of transmission rates in calendar time.

In some diseases, such as TB, there is considerable variation in the latency period, during which the mutation rate may be lower than it is during active disease. This variability can be incorporated into the negative binomial model as expressed in equation (14). We do not explicitly model within-host diversity, though this is relevant to identifying direct transmission events (Worby *et al.* 2014; Hall *et al.* 2015, 2016; Didelot *et al.* 2014, 2017). Cases of direct transmission will be clustered together with high probability in our method despite slight inaccuracy in the timing due to both branches of the pair’s two-case tree spending time in the same host. Pairs of cases for which the clustering decision is ambiguous are likely to have several intermediate cases between them, with a larger tree height, and so the contribution of in-host diversity in either sampled case will be small. In-host diversity in unsampled cases would not affect our estimates unless it contributed to changes in the molecular clock rate.

WGS data has been noted to be helpful in ruling out transmission but insufficient, on its own, to resolve transmission events (Casali *et al.* 2016; Campbell *et al.* 2018). If the primary use of WGS data is only to refute transmission, one might ask why clustering matters. We would argue that the transmissions that are not refuted by WGS are then presumably considered to be possible recent, or direct, or clustered transmissions. Even if the primary use of WGS data is to refute direct transmission, there is a trade-off between the strength of that refutation and the possibility of mistakenly refuting genuine recent transmission events. This is more likely, using SNP cut-offs, where selection (say for antibiotic resistance) has led to higher SNP differences than expected. In addition, in practice WGS data are not only used to refute direct transmission, but to produce clusters that inform onward analyses, reports on the extent of recent transmission, outbreak analysis and reconstruction and even public health policy; see (Guthrie *et al.* 2018) for one example.

We have accommodated the possibility of low substitution rates in latency with a non-Poisson model for the clock process, *λ*, in Equation (5) (though we have not implemented this) and to some extent with the option of a non-constant transmission rate. However, we have not modelled the possibility of a direct relationship between low SNP accumulation and low probability transmission. If this relationship exists - for example if latent cases both do not transmit and do not accumulate SNPs (Colangeli *et al.* 2014) – then low SNP differences could correspond to fewer intermediate hosts despite long elapsed times. This is an implicit assumption of a SNP-only method; while it may be correct it is a strong assumption, and there is evidence that mutation rates in latency are not reduced compared to active disease (Ford *et al.* 2011; Lillebaek *et al.* 2016).

We have not used the probability of sampling in forming our clusters, in contrast to other tools including the vimes package (Jombart and Cori 2017). For example, if it is known that surveillance is strong, then it would be less likely for 10 intermediate cases to be unsampled than for 5 intermediate cases, and this could be built in to a clustering method. Our rationale for not taking this into account is to provide a clustering approach that is as parallel as possible to the SNP cut-offs currently in widespread use while taking additional information on timing, molecular evolution and transmission into account. It is often the case that the true sampling rate is not known and may change over time, and – particularly for TB in high-resource settings – cases can be missed because they are hard to identify (perhaps being at higher risk of TB due to homelessness or other factors, as in Casali *et al.* (2016)). In many settings the sampling probability may be uncertain. We have taken the approach of defining the clusters themselves without explicit reference to the sampling probability, with the view that the clusters are central inputs to other analyses which will take sampling into account (as is done for example in TransPhylo (Didelot *et al.* 2017)). However, in our approach, changes in the sampling probability would likely be apparent in changes in the temporal and genetic distance between cases over time.

We have also not modelled changes in the transmission process over time in a community (eg due to depletion of susceptible individuals, improved infection control, etc). As with including sampling, this may best be done in a more nuanced analysis after the initial clustering rather than as part of the clustering itself, but in principle, changes to the transmission function over calendar time could be incorporated into the mathematics behind Equation (8). However, this would raise interpretation challenges because of the fact that our transmission process reflects the rate of the pathogen moving between hosts where it is known that there is an infected host at the "end" of the chain (since each pair consists of two sampled hosts, whose pathogen was sequenced and who were therefore certainly infected). We do not model the number of contacts over which transmission could have occurred.

The choice of a particular SNP cut-off also takes no account of the inevitable uncertainties involved in the gathering and processing of raw read data, and does not allow for the modelling of this uncertainty. Different bioinformatics pipelines - and different parameters used within those pipelines - can have a substantial effect on the number of SNP differences reported between cases. It is usual for SNP differences to be taken as given and, although sometimes details are provided – see for example Katz *et al.* (2013) - it is important to recognise that there can be considerable variation between SNPs reported using different pipelines and parameters. For example, the level of quality scores and read depth cut-offs used will generally have a high impact, as will the precise way in which hyper-variable sites and repeat regions are handled (or excluded). As technology improves we may begin to capture variation in repeat regions, or types of variation (eg insertions/deletions) that are currently masked, and in that new pipeline 12 SNPs may not carry the interpretation it does today. The model could easily incorporate more genomic information, resulting in a more sophisticated version of the distance function. In particular, large-scale genomic features can readily help to establish that cases belong to separate and therefore distantly related lineages. As variation-calling pipelines evolve, our method could be used to relate each pipeline to numbers of transmissions or to estimated divergence time; this would form an approach to compare bioinformatics pipelines and data sources, and to curate their use in defining distances between isolates.

TB has distinct phylogeographic lineages which have been reported to have different mutation rates, with lineage 2 (the East Asian and Beijing lineage) having higher mutation rates than lineage 4 (Euro-American) (Ford *et al.* 2013). Our approach could unify clustering despite such differences, as the same transmission and probability settings could be used under different SNP accumulation rates. This would provide a consistent approach to clustering in areas where multiple lineages co-circulate, and allow comparison of TB clustering patterns in different settings. The same would be true for adapting to differing natural histories across different pathogen lineages or sub-populations: the choice of *β* could reflect transmission differences while the other settings remained the same.

The long-term aim of changing how cases are assigned to clusters is to improve the way that WGS and epidemiological data are used and to best capture clusters that correspond to transmissions of an infectious disease. We have found that basing clusters on the number of transmission events, with a probabilistic cut-off, is feasible, can integrate timing and other data, and compares favourably to clustering based on SNP cut-offs.

## Materials and Methods

### Data

In this paper we focus on TB, but our approach is applicable to other pathogens for which whole genome sequencing can be carried out and where it is appropriate to use SNPs to compare closely-related isolates (naturally, parameters will vary). TB provides a convenient model as it avoids the complications associated with horizontal gene transfer, it is an important pathogen worldwide, it has very diverse epidemiological settings and WGS tools are increasingly used for public health purposes.

#### British Columbia

The British Columbia Centre for Disease Control (BCCDC)’s Public Health Laboratory (BCPHL) receives all *Mycobacterium tuberculosis* (Mtb) cultures for the province and performs routine MIRU-VNTR genotyping on all Mtb isolates. Mtb isolates belonging to MIRU-VNTR cluster MClust-012 were revived from archived stocks, DNA extracted, and sequenced using 125bp pairedend reads on the Illumina HiSeqX platform at the Michael Smith Genome Sciences Centre (Vancouver, BC). The resulting fastq files were analyzed using a pipeline developed by Oxford University and Public Health England. Reads were aligned to the Mtb H37Rv reference genome (GenBank ID: NC000962.2), with an average of 92% of the reference genome covered. Single nucleotide variants (SNVs) were identified across all mapped non-repetitive sites. Fastq files for all genomes are available at NCBI under BioProject PRJNA413593.

#### Republic of Moldova

##### Sample collection and epidemiological data

The study population included patients diagnosed with culture positive tuberculosis at the Municipal hospital from October 2013 - December 2014 in the Republic of Moldova. All epidemiological and laboratory data from TB patients are routinely entered into a country-wide web-based TB electronic medical record (EMR) database. Epidemiological data including age, sex, previous TB history, results of chest radiograph, history of incarceration, and place of residence were collected. Laboratory data, including mycobacterial smear grade, culture and drug-susceptibility testing to first and second line anti-tuberculosis agents, were extracted from the EMR. As part of this study, all *M. tuberculosis* patient isolates were subcultured and frozen for genomic analysis.

##### Variant calling and phylogenetic analysis

DNA was extracted from M. tuberculosis grown on Lowenstein-Jensen slants as described previously. Paired-end (250 base pair) sequences were generated on the Illumina MiSeq platform. Raw fastq reads were filtered for length and trimmed for low-quality trailing base pairs using Trim Galore, aligned to the H37Rv NC000962.3 reference genome using BWA, with duplicate reads removed using PicardTools. The mpileup function in samtools was used for single-isolate variant calling. Isolates with a high proportion of apparent mixed or heterozygous single nucleotide polymorphism (SNP) calls (i.e. those with >25% reads supporting the reference allele) were excluded from analysis. SNPs within 15 base pairs of insertions or deletions (indels) or with variant quality scores < 100 were excluded. SNPs in or within 50 base pairs of hypervariable PPE/PE gene families, repeat regions, and mobile elements were excluded (Eldholm *et al.* 2015). A phylogenetic tree was constructed in RAxML (GTR-gamma for nucleotide substitution and correcting for SNP ascertainment bias) and annotated with DST results and drug-resistance associated variants from Mykrobe Predictor (Bradley *et al.* 2015). Representative strains from other studies in the region, including L4 (LAM, Haarlem, Ural) and L2 (Casali *et al.* 2014; Merker *et al.* 2015), were also included. Percy256 (Lineage 7) was included as an outgroup. Fastq files for all genomes are available at NCBI under Accession number SRP156366.

### Clustering approach

The overall approach is to use the SNPs and case timing to derive a distribution for the time to the MRCA of each pair of samples, condition on that time to write the probability that the samples were separated by some number of transmission events, and then integrate out the unknown time to the pairs’ MRCA. The first step makes use of the molecular clock process and depends on the clock rate and on the numbers of SNPs under a form of selection (like antibiotic resistance). The second step using information about the transmission process and the natural history of the pathogen.

For each sample, we start with the date on which the sample was taken and the aligned nucleotide sequence for the set of variable sites in our set of samples. For any two samples *S*_1_ and *S*_2_, we have the SNP distance *N* = *N*(*S*_1_, *S*_2_) which is equal to the Hamming distance between their respective nucleotide sequences. We also have the sampling time difference *δ* = *δ*(*S*_1_, *S*_2_). Without loss of generality, we can assume that *S*_1_ is sampled either at the same time as, or before, *S*_2_. What we do not know *a priori*, and therefore we have to estimate, is the total amount of time *h* over which the SNPs have accumulated (on both branches in total) since the date of the MCRA of *S*_1_ and *S*_2_. We also refer to *h* as the “height”.

Given the time *h*, we can use a transmission process to estimate the probability that there are more than some threshold number of transmission events *T* in a total time *h*; we integrate over the unknown *h*. This transmission process need not be homogeneous.

We make various assumptions in setting up the model. Both substitutions and transmissions occur according to (possibly non-homogeneous) Poisson processes over time. Unless it is otherwise stated, the population from which the samples are drawn is homogeneous, so transmission is random and equally likely between hosts irrespective of factors such as location of abode, individual lifestyle etc. We do not assume that all infected cases are reported and sequenced. However, where we do have sequence data, we assume that it is correct and complete. Reported cases may be sampled more than once. We do not explicitly model reporting and sampling rates. If these change over time, then this would be reflected in the time and genetic distance between nearby cases and consequently in the estimated number of intermediate transmissions between reported cases. Once infected, we assume that a patient becomes infectious immediately, either with a constant probability of infection per unit of time, or in a process yielding a gamma-distributed time to the next infection. This "natural history" model is assumed not to change with calendar time, such that the course of infectiousness proceeds in the same manner from infection to infecting others independent of the calendar time of infection. Our approach is intended to group sequences into clusters, and does not model reported cases for whom there is no sequence data.

Noting that *δ* is fixed by the sampling times in the data, we estimate the distribution of the time *h*/2 over which the SNPs have had to accumulate before the sample date of *S*_1_. This is equivalent to estimating the date of the MRCA of *S*_1_ and *S*_2_. Because both branches are free to evolve over this time, *h*/2 + *h*/2 = *h* is the effective overall time between the MRCA and *S*_1_, and *δ* + *h* is therefore the total evolutionary time separating the two cases.

#### Estimate of the height where sample dates are the same

The simplest model for the number of SNPs per unit time is a Poisson process with a constant rate *λ*; we can also accommodate overdispersion, reflecting a more variable SNP accumulation process suitable for pathogens whose substitutions are not as clock-like (see below). The standard Poisson distribution with parameter *λh* gives the probability density of the number of SNPs on a given time interval *h*:

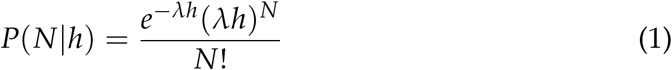

However we are interested in the likelihood of the time *h* as a function of the specified number of SNPs.

We know by standard theory that the arrival time density – that is, the time density until the next SNP – can be modelled by the exponential density function *λe*^−*λh*^. Furthermore, the waiting time until the *N*-th SNP is also a Poisson process, as the arrivals are assumed to be independent and identically Poisson distributed. It can be shown (for example in Chapter 2 of (Gallagher 2013)) by repeated convolution of densities that the distribution of the *N*th arrival time *A*_*N*_ is given by

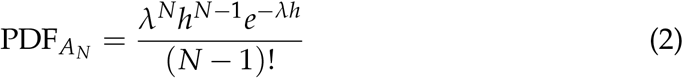

for *N* > 0. This is the Erlang distribution, with mean = *N*/*λ*, as expected.

We know that exactly *N* SNPs have already occurred on a time interval of uncertain length *h*, and we are interested in the likelihood of *h* given the data *N*. Since we already have *N* SNPs and are waiting for the (*N* + 1)-th, this is given by the arrival time density for the (*N* + 1)-th SNP; by replacing *N* with *N* + 1 in the above and interpreting it as a function of *h*, we have:

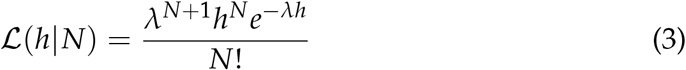

Note that when *N* = 0, this reduces to *λe*^−*λh*^.

Alternatively, we can generalise the arrival time density to a gamma distribution, where the extra parameter allows us to fix the mean but change the variance. This allows us to be more flexible with respect to dispersion than with using the exponential distribution. The gamma density, with two parameters *a* and *b*, is

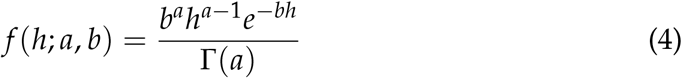

The mean is *a*/*b* and the variance *a*/*b*^2^. Note that we can recover the Poisson model result by setting *a* = 1 and *b* = *λ* (Cameron and Trivedi 2013). In this case the arrival time density for the (*N* + 1)-th SNP is given by

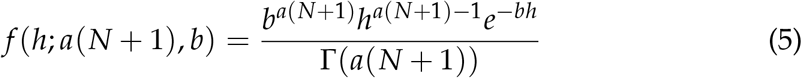

by standard properties of the gamma distribution.

#### Estimate of the height where sampling times differ

In this case, we account for the fact that some of the SNPs may have occured in the the fixed time interval of length *δ* between the two sample dates. Again, we begin with the simple model in which the number of SNPs occurring in this time is given by a Poisson distribution, in this case with parameter *λδ*. We write *N* = *N*_*h*_ + *N*_*δ*_, where *N*_*δ*_ is Poisson distributed with parameter *λδ*.

The number of SNPs *N*_*δ*_ accumulated on the fixed interval of length *δ* is somewhere between 0 and N inclusive; 0 ≤ *N*_*δ*_ ≤ *N*. Unconstrained, *N*_*δ*_ is Poisson distributed with parameter *λδ*. Conditioning on the probability that *N*_*δ*_ does not exceed *N* gives us the probability density

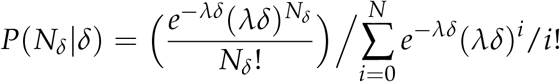

and writing *F*(*N*_*δ*_) for 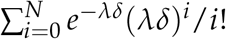

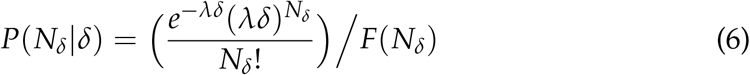

To obtain the expression for 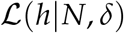, we sum over all the possible values of *N*_*δ*_, giving

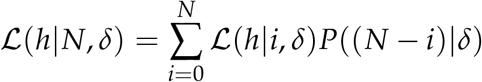

Substituting into our earlier expression,

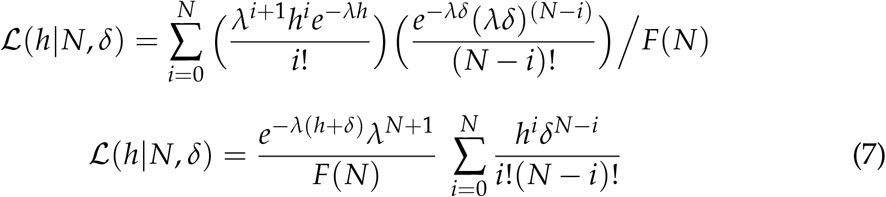

An example plot for the equation above is shown in Figure 5.

**Figure 5:**
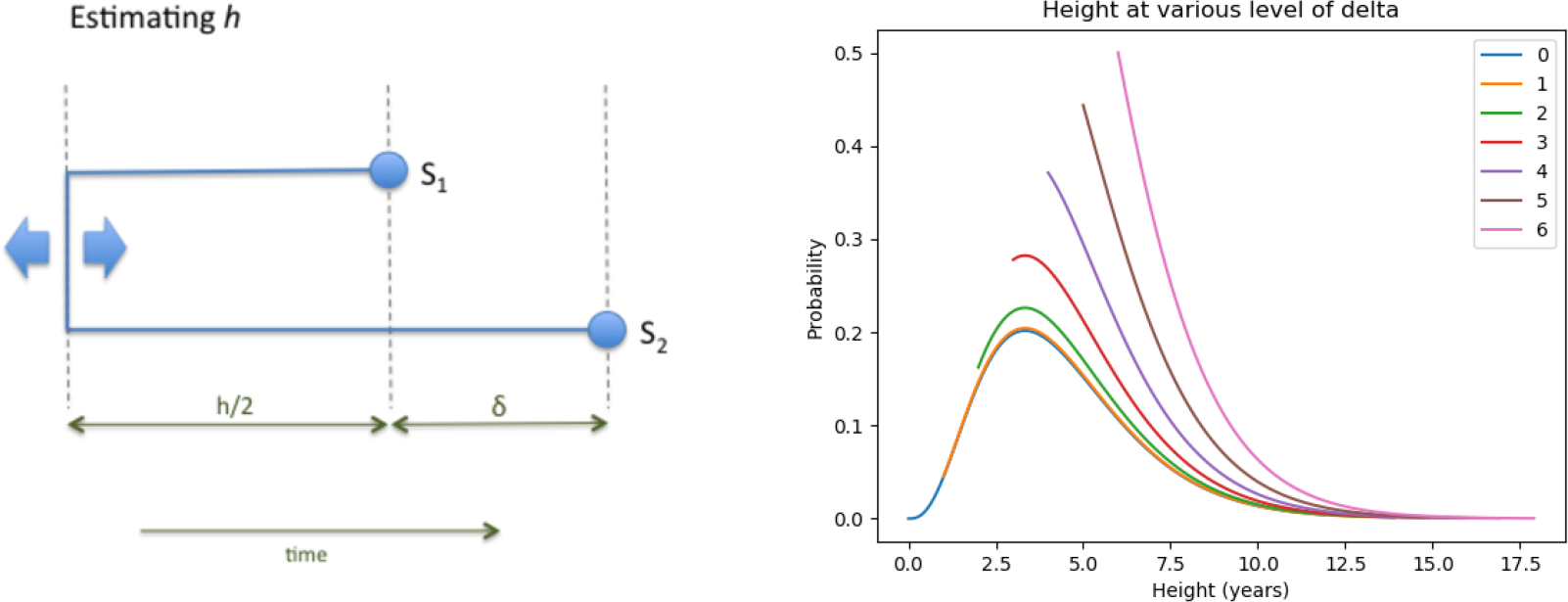
On the left is a schematic illustration of the notation. *h* is the total time in years over which SNPs accumulate between two cases before the first sample is taken, whereas the total time over which SNPs can occur is *h* + *δ* years. *δ* is known and fixed; *h* is unknown. On the right is a plot of *h* + *δ*, where *h* is given by Equation (4), for values of *δ* ranging from 0 through 6 years, with *N* = 3, *λ* = 0.9 SNPs/genome/year, and *β* = 1.2 transmissions/year. Since *h* + *δ* > *δ*, the lines corresponding to higher values of *δ* begin above 0.

#### Modelling transmissions

We connect SNP distances to transmissions using a model for the number of transmissions likely to have occurred over a given total time period, *conditional* on the two cases being infected at or before the sampling times. This means that unlike a transmission rate in a population-level epidemic model, which typically describes the rate of transmission per unit time given contact between a susceptible and an infectious individual, our transmission process is better described in terms of the rate at which a pathogen lineage will jump to a new host. This is, of course, distinct from the rate at which new transmissions occur in a community and the per-contact rate of transmission of infection between two individuals. We first assume for simplicity that *β* is a constant function, and that it is a Poisson process; we allow a more general model later. The amount of time over which transmissions can occur between our two cases is *h* + *δ*, and the expected number of transmissions is *β*(*h* + *δ*). The number of transmission events *k* is therefore given by

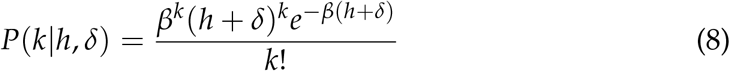

Integrating over *h*, we have

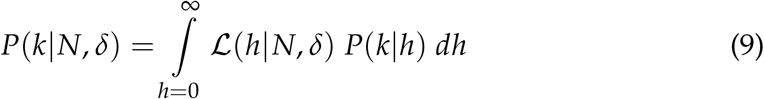

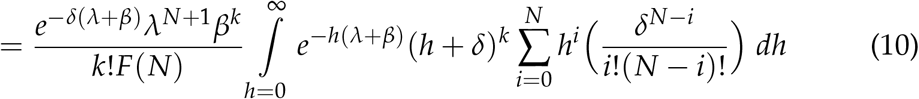

This equation expresses the key relationship that allows us to translate raw SNP differences and sample time differences into transmission probability distributions - examples are shown in Figure 6. As the sample time between cases increases, it can be seen that this factor makes an increasingly important contribution, relative to the SNP distance, to the distance between cases.

**Figure 6:**
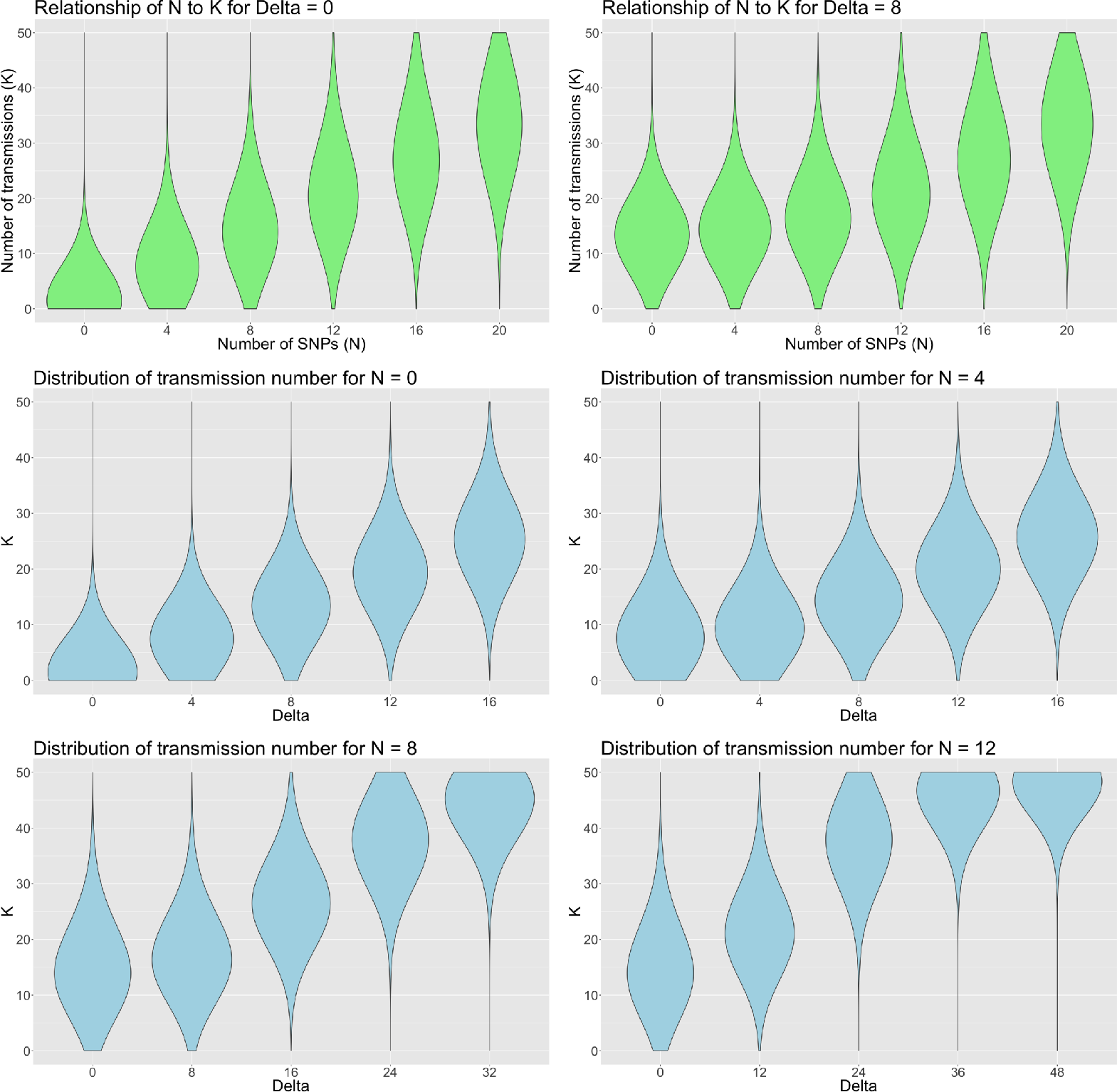
Probability density for the number of transmissions, given by Equation (10), with clock rate *λ* = 0.9 SNPs/genome/year and *β* = 1.5 transmissions/year. The upper two panels show the densities for delta values of 0 and 4 years for a range of SNP distance between 0 and 20. The lower four panels show the densities for 0, 4, 8 and 12 SNP distance respectively, for *δ* = 0, 4, 8, 12, 16 years.

Unless stated otherwise, Equation (10) is used to generate the data presented in the Results Section.

#### Time varying transmissions

In our context, a transmission event should be understood as an event in which a pathogen is transferred to a new host, ultimately causing a secondary case in that host. While there may be undetected transmission events in which the secondary cases never develop disease, our data are on sampled cases with active disease, and the time between successive transmissions should approximately reflect the serial interval between cases with active disease. We allow the number of transmissions *β* = *β*(*t*) to be a function of time since infection, allowing for a variable risk of infecting others during the course of infection. Once a host is infected, the details of the natural history of the pathogen affect the generation time - the exact form of the function *β*(*t*) allows us to incorporate the varying rates of progression from infection to active disease, and then on to transmission. This illustrates that our framework has the flexibility to include more detailed and accurate modelling of the underlying disease dynamics. As stated in Didelot *et al.* (2017), the generation time distribution can take any form (Fine 2003; Wallinga and Lipsitch 2007), but gamma distributions are often used, as for example in Conlan *et al.* (2010). We apply the gamma distribution with parameters shape *α* and scale *θ*, so that:

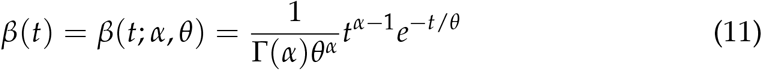

and the mean value is *αθ* * *dt* in a given time interval *dt*.

Putting this together with our Poisson model for the number of SNPs on a time interval (Equation (1)), we obtain:

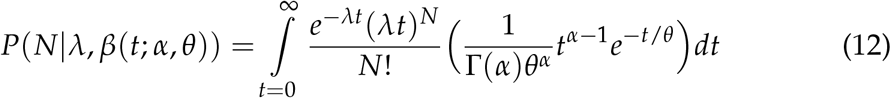

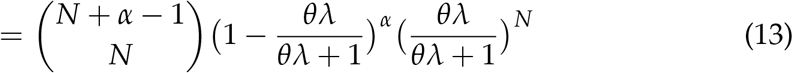

This is a negative binomial (denoted NB) distribution for the number of SNPs for one transmission generation,

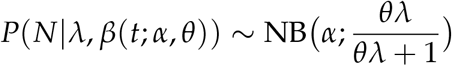

Assuming transmission events are independent of each other, it then follows (by standard properties of the negative binomial) that the probability of *N* given *k* transmissions is also distributed as a negative binomial, with

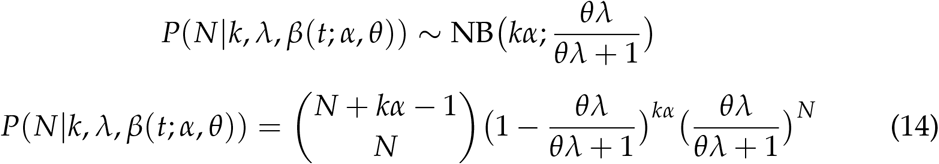

#### Modelling resistance-conferring SNPs

Suppose that we know that there are resistance-conferring SNPs in our sample population, or perhaps other SNPs at sites known to be under selection or simply to have a different rate of substitution. Let us assume they account for a certain fixed proportion of the observed SNP differences. Given *N* SNPs, assume that *m* are not resistance-conferring and *n* are, so *N* = *m* + *n*. Their respective mutation rates are given by *λ*_*m*_ and *λ*_*n*_, where *λ*_*n*_ > *λ*_*m*_. Assuming independence, on a given time interval of length *h* we have

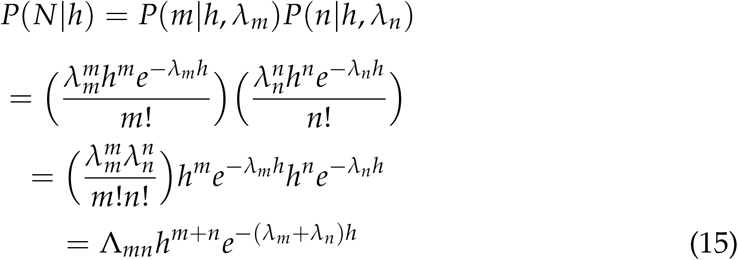

where

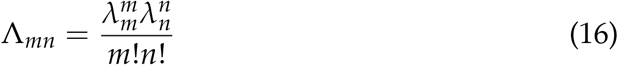

Compare this to Equation (1), which we can recover by setting *m* = *N*, *n* = 0, *λ* = *λ*_*m*_, and *λ*_*n*_ = 0.

We have a Poisson process which is the sum of two independent Poisson processes with *λ* = *λ*_*m*_ + *λ*_*n*_. As before, we can derive expressions for ℒ(*h*|*N* = *n* + *m*) and *P*(*k*|*N* = *n* + *m*), so

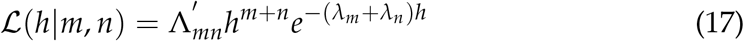

where

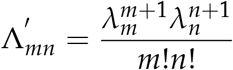

A way to illustrate the effect of including resistance-conferring SNPs is to consider the expected value of *h*. Recall that under Equation (1), the mean is given by *N*/*λ*. Thinking of our resistance and non-resistance-conferring SNP processes, they have means respectively of *n*/*λ*_*n*_ and *m*/*λ*_*m*_. Thus the combined process has mean *n*/*λ*_*n*_ + *m*/*λ*_*m*_, and we can write the rate parameter *λ*\* of the combined process as

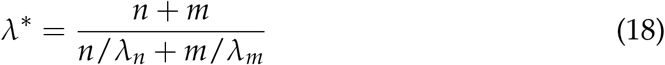

Note that for large *λ*_*m*_, *λ*\* tends to *λ*_*n*_ * (*n* + *m*)/*n*. The larger the value of *λ*_*m*_ as compared to *λ*_*n*_, the smaller the contribution that the resistance-conferring SNPs make to the value of *h* - accordingly, 4 SNPs likely to have arisen due to inappropriate treatment or another selection process should not contribute as strongly towards separating two cases into different transmission clusters as 4 "neutral" SNPs. Ideally, the value of *λ*_*m*_ should be estimated from data. Once resistance SNPs have occurred in an individual, they are likely to be transmitted onwards when the individual infects others. These secondary cases share the resistance SNPs with each other (*n* = 0 in these pairs) and they are likely to be placed in the same cluster. Between each secondary case and the infecting case, *n* > 0; our method allows the resistance SNPs to "count for" less time than other SNPs, and the index case is likely clustered with the onward cases.

#### Spatial proximity and other individual data

Other factors that affect the likelihood of transmission, such as spatial proximity or other covariate data including contact tracing, demographics or other host factors, can be built into the model.

To incorporate spatial proximity, we assign each of the cases into one of a number of regions *R*_*i*_ where *i* is the region index. For the British Columbia data set, there are six regions defined, as shown in Figure 3. For any pair of cases, a probability weighting is assigned which is equal to 1 in the case that both cases belong to the same region, and a value below 1 for cases which belong to different regions. This weighting *w* is then applied to the probability of obtaining *k* transmissions given *N* SNPs, giving us a modified version of Equation (10)

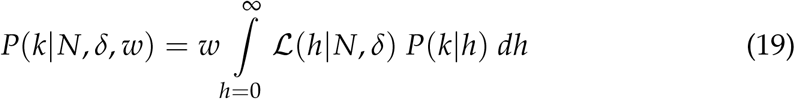

#### Simulations

We generate simulated outbreaks and compare the SNP and transmission methods on them with a technique that measures the similarity of clusters using an information-theoretic approach (Meilă 2007). Outbreaks are simulated using *TransPhylo* (Didelot *et al.* 2017), which generates a dated transmission network for each simulation, containing both sampled and unsampled cases. From these, and for all the cases, phylogenetic trees are extracted using *phyloTop* (Kendall *et al.* 2016). Sequences are then generated with *phangorn* (Schliep 2011) and output as *fasta* format files. For the sampled, and therefore "known", cases we generate sets of clusters using the SNP and transmission methods for a range of cut-off levels. We also generate the "true" clustering of the sampled cases implied by the simulated *TransPhylo* transmission networks.

### Software availability

The methods presented here are available as R functions in the `transcluster` package, available at https://github.com/JamesStimson/transcluster.

## Supporting information

## Acknowledgements

CC and JS are supported by EPSRC grant EP/K026003/1 and CC is additionally supported by EPSRC grant EP/N014529/1. TC is supported in part by U54GM088558 from the National Institute of General Medical Sciences. The content is solely the responsibility of the authors and does not necessarily represent the official views of the National Institute Of General Medical Sciences or the National Institutes of Health. We would like to thank Tyler S. Brown for contributions to the bioinformatics and analysis of the data from Moldova.

