## Supplementary material for "Beyond the SNP threshold: identifying outbreak clusters using inferred transmissions"

December 3, 2018

### Application of transmission method to timed trees

We can extend the transmission method by applying it to timed phylogenetic trees. Building such a tree (using, for example, Beast2 (Bouckaert *et al.* 2014)) allows us to consider the joint ancestry of all isolates together, in contrast to the pairwise application in the main text. Tree reconstruction algorithms account for varying mutation rates at different sites (which can be specified or estimated), incorporate evolutionary models that discriminate between transitions and transversions, account for various population models and have other flexibilities. In Bayesian tree reconstruction, the timings of the branches are obtained from the posterior, and timing and sequence data are jointly used to construct a phylogenetic tree in which the branch lengths are in units of time. An advantage of this approach is that the clock rate can be estimated from the data, rather than being a fixed assumption or a range, though naturally this requires longitudinal data and sufficient genetic variation.

To cluster isolates using a timed tree, the timed tree is subdivided into a set of sub-trees by removing internal branches that exceed the transmission cut-off. The cut-off length is obtained using Equation (8) in the Methods section of the main text, which gives the probability of transmissions based on the total time between nodes. In this case, we are considering the time between two internal nodes rather than the total time between two sampled cases (which are tips of the tree), so we replace $h + \delta$ with the branch length between any two internal nodes. Note that we do not cut terminal branches. Cutting a branch results in a sub-tree being created from the clade descended from this branch. In the original tree, the cut branch and its descendant clade are then replaced by a single terminal branch.

In Figure S1 we illustrate the application of this method for a simulated data set containing 22 samples taken over a 10 year period. Note that not all of the clusters obtained are monophyletic clades. Cluster 2 and Cluster 3 are clades; but Cluster 1 is not, Cluster 2 being its phylogenetic descendant. This is a feature which is also obtained by Barido-Sottani *et al.* (2018) in the context of HIV transmission clusters, based on a multi-state birth-death model with variable transmission rates.

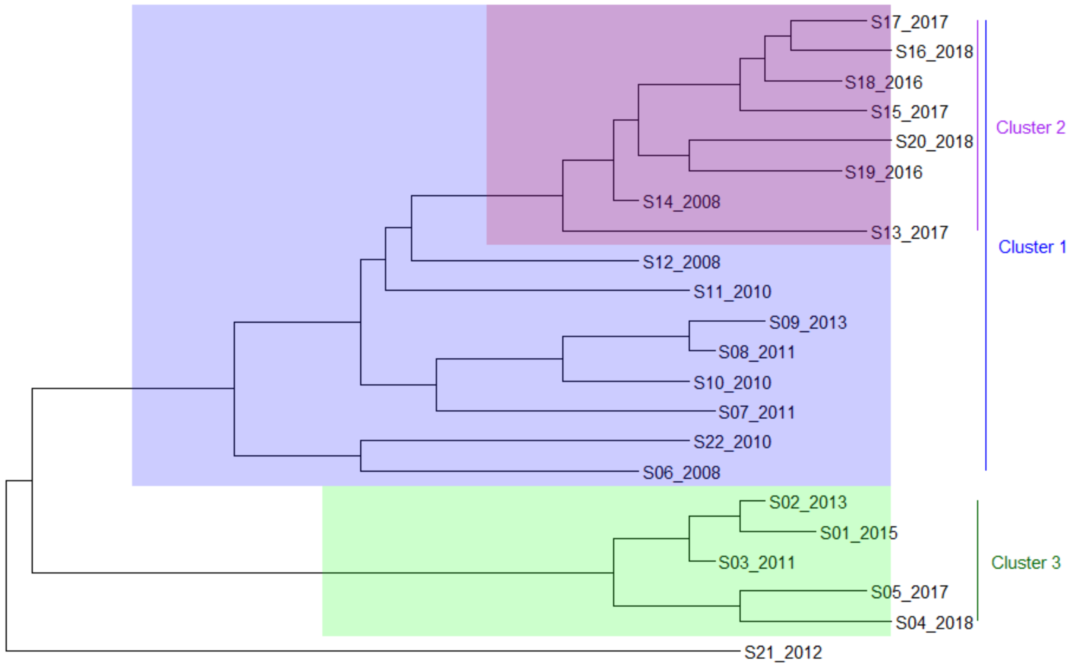


Figure S1: Application of transmission method to a simulated timed tree. Branches are cut where the shading colour changes, which is where there is a greater than 80% probability that at least 10 transmissions have occurred along that branch, with $\beta = 2.2$ transmissions/year. This occurs for any internal branch with length at or in excess of 6 years. Three sub-trees are created, corresponding to the identified clusters 1, 2 and 3. Tip labels are suffixed with the year of sampling.

The function *clusterTimedTree*, available in the R package `transcluster`, was used to partition the example in Figure S1.
